# Structural alterations in the macaque frontoparietal white matter network after recovery from prefrontal cortex lesions

**DOI:** 10.1101/2020.05.02.064840

**Authors:** Ramina Adam, David J. Schaeffer, Kevin Johnston, Ravi S. Menon, Stefan Everling

## Abstract

Unilateral damage to the frontoparietal network typically impairs saccade target selection within the contralesional visual hemifield. Severity of deficits and the degree of recovery have been associated with widespread network dysfunction, yet it is not clear how these behavioural and functional changes relate with the underlying structural white matter pathways. Here, we investigated whether recovery after unilateral prefrontal cortex (PFC) lesions was associated with structural white matter remodeling in the distributed frontoparietal network. Diffusion-weighted MRI was acquired in four macaque monkeys before the lesions and at 2-4 months post-lesion, after recovery of deficits in saccade selection of contralesional targets. Probabilistic tractography was used to reconstruct inter- and intra-hemispheric frontoparietal fiber tracts: bilateral superior longitudinal fasciculus (SLF) and transcallosal fibers connecting bilateral PFC or bilateral posterior parietal cortex (PPC). After behavioural recovery, tract-specific fractional anisotropy in contralesional SLF and transcallosal PPC increased after small lesions and decreased after larger lesions compared to pre-lesion. These findings indicate that remote fiber tracts may provide optimal compensation after small PFC lesions. However, larger lesions may have induced widespread structural damage and hindered compensatory remodeling in the structural frontoparietal network.

## 1. Introduction

Impaired spatial attention and reduced gaze shifts toward the contralesional visual hemifield are commonly seen following unilateral damage to the primate frontoparietal network, which includes the caudal prefrontal cortex (PFC), posterior parietal cortex (PPC), and white matter pathways connecting the large-scale network (Bartolomeo et al., 2012; Corbetta and Shulman, 2011; Mesulam, 1999). In stroke patients, these deficits manifest as a decreased ability to respond or attend to a single visual target within the contralesional hemifield, a phenomenon known as visual neglect (Bartolomeo, 2007; Li and Malhotra, 2015). In many cases, deficits within the contralesional hemifield appear only in the presence of a competing stimulus in the ipsilesional hemifield, referred to as visual extinction (Bisiach, 1991; Damasio et al., 1980; B. de Haan et al., 2012; Di Pellegrino et al., 1997). Similar visuospatial deficits within the contralesional hemifield have been demonstrated in macaque monkeys after experimental lesions or reversible inactivation of PFC or PPC areas (Adam et al., 2019; Bianchi, 1895; Deuel and Farrar, 1993; Johnston et al., 2016; Latto and Cowey, 1971; Lynch and Mclaren, 1989; Rizzolatti et al., 1983; Schiller and Chou, 1998; Wardak et al., 2002, 2006, 2004; Wilke et al., 2012). Functional imaging studies of visual neglect and extinction in patients and animal models have shown that functional changes across a distributed network correlated with the severity of deficits in the acute stage (Baldassarre et al., 2014; Umarova et al., 2011; Wilke et al., 2012) and with the degree of behavioural recovery in the chronic stage (Deuel and Collins, 1983; He et al., 2007; Ramsey et al., 2016; Umarova et al., 2016).

Recently, we reported the longitudinal changes in resting-state functional connectivity (rsFC) within the frontoparietal network after a unilateral caudal PFC lesion in macaque monkeys (Adam et al., 2020). We showed that recovery of contralesional saccade choice deficits correlated with increasing rsFC between the contralesional PFC and ipsilesional PPC. Since network-wide rsFC has been shown to reflect properties of the underlying structural connections (Dijkhuizen et al., 2012; Greicius et al., 2009; Hagmann et al., 2008; Shen et al., 2015), here we expand on our previous study to examine the lesion-induced structural changes of white matter pathways connecting the bilateral frontoparietal network, including bilateral superior longitudinal fasciculus (SLF) and transcallosal fibers connecting bilateral PFC and bilateral PPC. The SLF is long-range association pathway that connects frontoparietal areas within hemisphere (Petrides and Pandya, 1984; Schmahmann et al., 2007; Thiebaut de Schotten et al., 2012). Between hemispheres, the caudal PFC and PPC are connected to their respective contralateral homologs via transcallosal fibers which cross at the genu or isthmus of the corpus callosum, respectively (Barbas and Pandya, 1984; Hofer et al., 2008). It has not yet been explored whether recovery of contralesional target selection after a focal lesion is associated with changes in related white matter fibers. Moreover, the behavioural relevance of structural alterations in remote fiber tracts before and after focal damage have been understudied and are not well understood.

In the present study, we examined the microstructural changes of frontoparietal white matter tracts in those four macaque monkeys using high spatial and high angular resolution diffusion-weighted MR imaging (DWI) acquired *in vivo* at 7T. DWI data were obtained at two time points: before the unilateral PFC lesion and at a late post-lesion stage (week 8 or 16 post-lesion) when contralesional saccade choice deficits had largely recovered. Probabilistic fiber tractography was used to reconstruct four fiber tracts of interest: contralesional and ipsilesional SLF and transcallosal PFC and PPC tracts. Tract-specific diffusion parameters, including fractional anisotropy (FA) and mean, axial, and radial diffusivity, were then calculated for each tract and compared over time. We speculated that the remote fiber tracts (i.e., contralesional SLF and transcallosal PPC) may have mediated the increased rsFC between contralesional PFC and ipsilesional PPC that was found in our previous study (Adam et al., 2020), since those tracts provide an undamaged pathway which indirectly links the cortical regions together. On the other hand, ipsilesional SLF and transcallosal PFC fibers were likely damaged by anterograde/retrograde degeneration since they directly innervate the lesioned right caudal PFC (Thomalla et al., 2004; Werring et al., 2000). Thus, we hypothesized that the remote contralesional SLF and transcallosal PPC tracts play a compensatory role to support behavioural recovery post-lesion, by potentially mediating increased rsFC of the frontoparietal network. We predicted that if behaviour or rsFC relied on the contralesional SLF and transcallosal PPC tracts, then FA should increase within one or both of those remote tracts from pre-lesion to late post-lesion.

## 2. Methods

### 2.1. Subjects

Data were collected from four adult male macaque monkeys (*Macaca mulatta*) aged 5 – 7 years old and weighing 7 – 10 kg. All surgical and experimental procedures were carried out in accordance with the Canadian Council of Animal Care policy on the use of laboratory monkeys and approved by the Animal Care Committee of the University of Western Ontario Council. Experimental methods for these subjects has been previously published in our companion paper (Adam et al., 2020). Herein, animals are individually described as Monkey L, Monkey S, Monkey B, and Monkey F. We show behavioural data from these subjects at the following time points: pre-lesion, week 1-2 post-lesion (early post-lesion), and week 8 or 16 post-lesion (late post-lesion). The early post-lesion time point shows the acute behavioural deficits following the lesion and the late post-lesion time point shows the recovered behaviour months later. DWI data was acquired at pre-lesion at late post-lesion (described below) to examine how the white matter microstructure changed at the time of behavioural recovery compared to pre-lesion.

### 2.2. Lesions

Details of the experimental lesioning surgeries have been previously published in these subjects (Adam et al., 2020). Briefly, lesions were induced using the vasoconstrictor endothelin-1, which induces focal occlusion with subsequent reperfusion and has been validated in marmosets and macaque monkeys (Dai et al., 2017; Herbert et al., 2015; Murata and Higo, 2016; Teo and Bourne, 2014). Intracortical injections were made in the right caudal PFC (along the anterior bank of the arcuate sulcus and the caudal portion of the principal sulcus). We varied the total amount of endothelin-1 administered to each animal to produce small lesions in Monkey L and Monkey S (6-12 μg) and large lesions in Monkey B and Monkey F (16-32 μg). Figure 1 shows the lesion extent in each animal. All monkeys sustained damage to the right caudal PFC with consistent lesions in area 8AD (Fig. 1B). The lesion in Monkey L was localized to area 8AD and 8B; in Monkey S, the lesion extended further into the dorsolateral and ventrolateral PFC. In Monkey B, the lesion extended into dorsal premotor areas and in Monkey F it extended into the dorsolateral and ventrolateral PFC and premotor areas. Lesion volume analysis showed that Monkey B and Monkey F sustained larger lesions than Monkey L and Monkey S with a lesion volume that was more than doubled (Monkey L = 0.43 cm^3^, Monkey S = 0.51 cm^3^, Monkey B = 1.28 cm^3^, Monkey F = 1.41 cm^3^).

**Figure 1.**
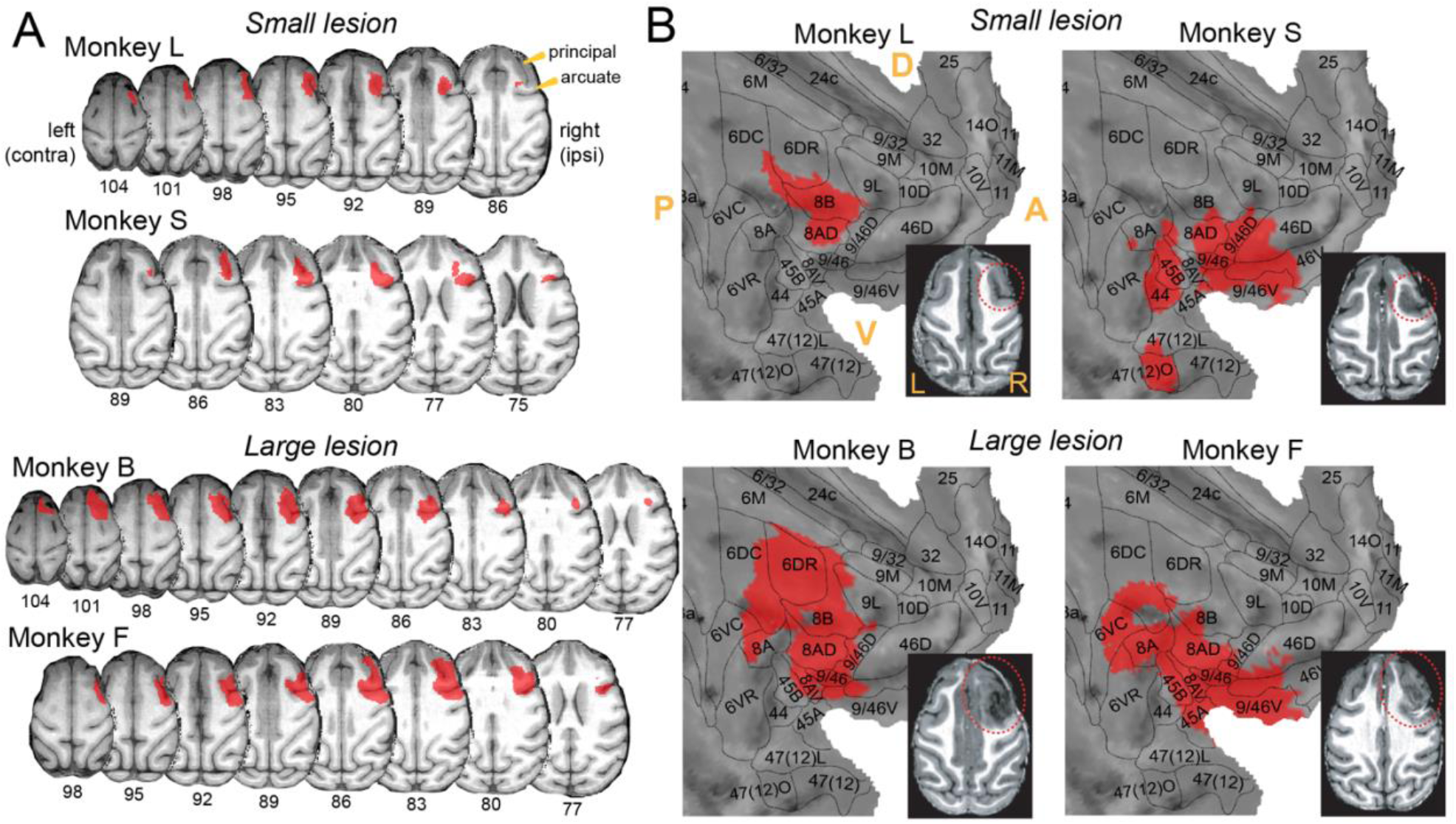
Lesion masks projected onto the macaque F99 template brain. Each monkey’s T1-weighted MP2RAGE anatomical image obtained one week post-lesion was segmented based on tissue type. Masks representing lesioned tissue were registered to the standard macaque F99 space and projected onto (A) axial slices of the macaque F99 brain and (B) cortical flat map representations of the macaque F99 right hemisphere with surface outlines that we created based on the Paxinos et al. (2000) macaque cortical parcellation scheme (Paxinos et al., 2000). Bottom right: one axial T1 image at one week post-lesion showing the lesioned tissue within the dotted red line boundary for each monkey. Abbreviations: principal = principal sulcus; arcuate = arcuate sulcus; contra = contralesional; ipsi = ipsilesional; D = dorsal; V = ventral; A = anterior; P = posterior; L = left; R = right.

### 2.3. Behaviour

We have previously reported the saccade target selection at pre-lesion and late post-lesion (Adam et al., 2019) but here we compare behavioural performance with DW-MRI data. For a detailed report of the behavioural task design, see (Adam et al., 2019). Before the lesion was induced, monkeys were trained on a saccade task that included two randomly interleaved trial types: (1) visually-guided single target trials and (2) free-choice paired stimulus trials in which a single visual stimulus appeared in each hemifield either simultaneously or at varying stimulus onset asynchronies (SOAs) and monkeys were able to freely select either stimulus as a saccade target to receive a liquid reward (Fig. 2). SOA is the variable temporal delay between presentation of the contralesional and ipsilesional stimulus on the free-choice trials. Each trial began with the presentation of a fixation point, followed by either a single visual target in the contralesional (left) or ipsilesional (right) hemifield or two peripheral stimuli, with one in the contralesional and one in the ipsilesional hemifield presented at a variable SOA. Free-choice trials were used to measure the degree of visual extinction, since contralesional extinction deficits only appear in the presence of a competing ipsilesional stimulus. This task is able to show whether a monkey exhibits a spatially lateralized saccade selection deficit by measuring saccade choice for contralesional and ipsilesional visual stimuli.

**Figure 2.**
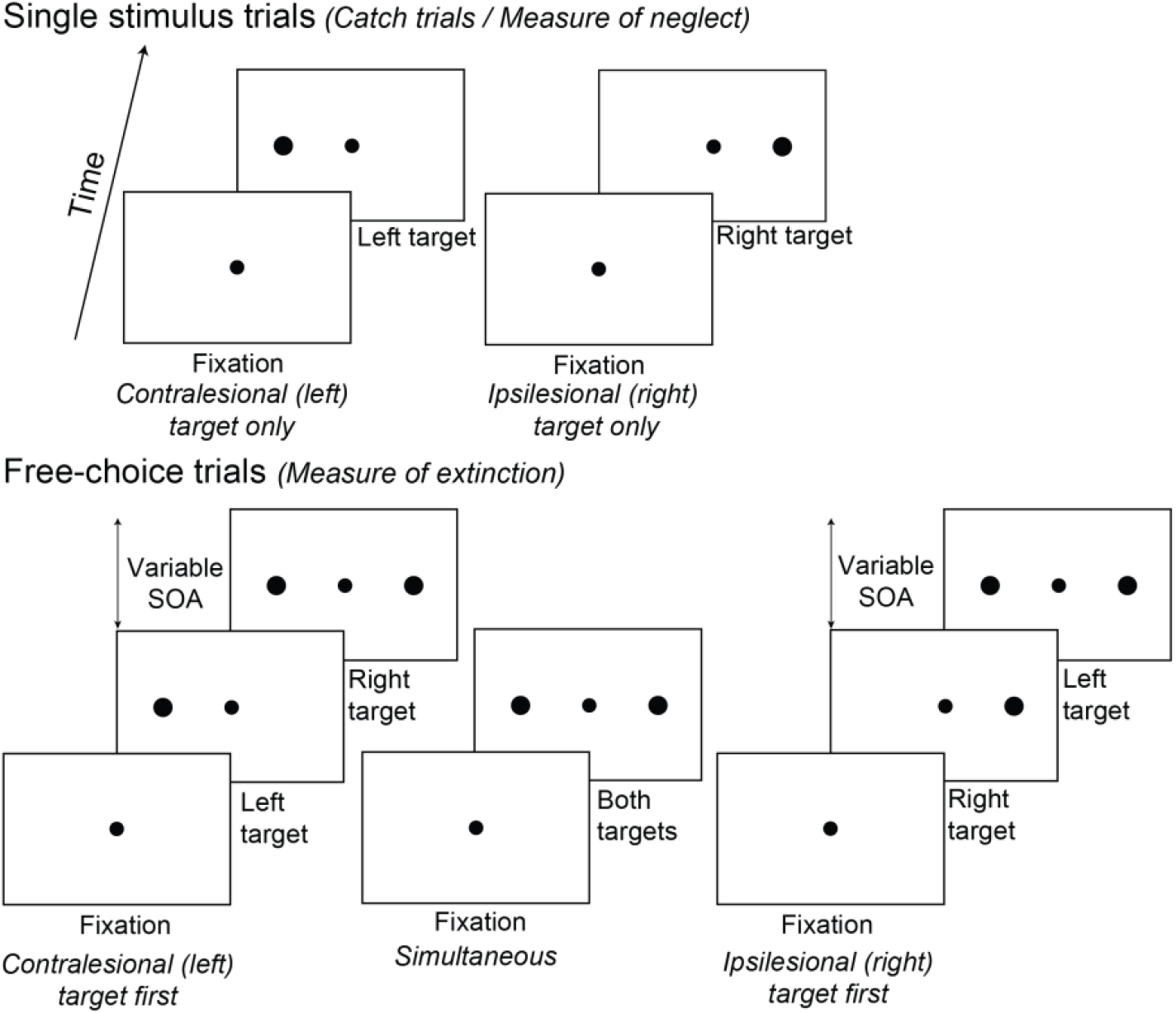
Behavioural task. Single target and free-choice paired stimulus trials were randomly interleaved within a session. Each trial began with the presentation of a fixation point, followed by either a single target in the contralesional (left) or ipsilesional (right) hemifield or two visual stimuli in either hemifield presented at a variable stimulus onset asynchrony. Stimulus onset asynchrony was the variable temporal delay (0-256 ms) between presentation of the left and right stimulus on the free-choice paired stimulus trials. Abbreviation: SOA = stimulus onset asynchrony.

In brief, we found that the right caudal PFC lesion induced deficits in contralesional target selection, such that there were decreased correct saccades made towards a single contralesional target (Fig. 3A) and decreased saccade choice of the contralesional stimulus on the free-choice trials (Fig. 3B,C). Deficits gradually recovered over 2-4 months post-lesion. We considered post-lesion behaviour as ‘compensated’ when task performance stabilized without further improvement. The time point for compensated behaviour was week 8 post-lesion for the small lesion monkeys (Monkeys L and S) and week 16 post-lesion for the large lesion monkeys (Monkeys B and F); we refer to these time points collectively as ‘late post-lesion’. Detailed results on this behavioural paradigm have been previously published (Adam et al., 2020, 2019).

**Figure 3.**
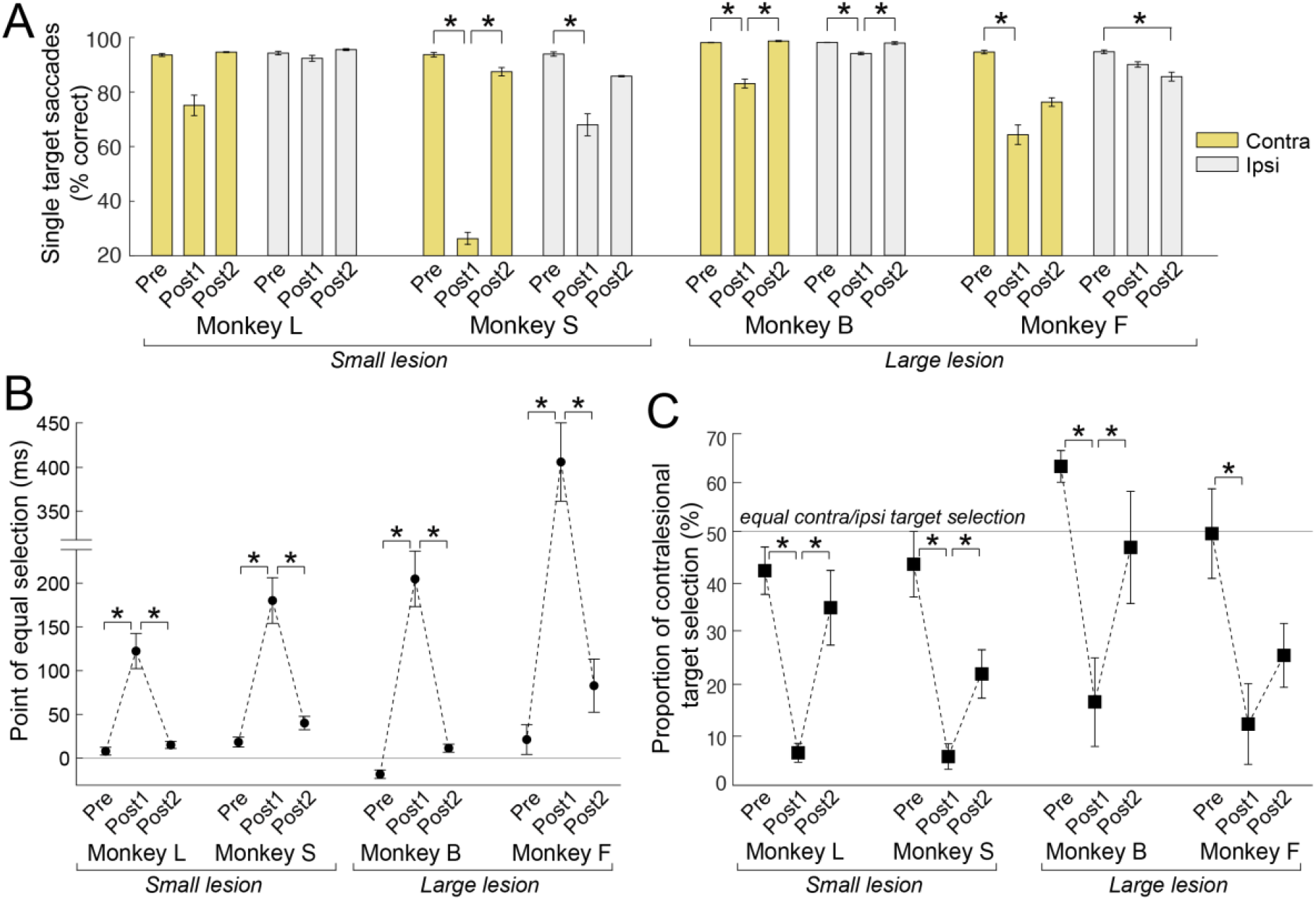
Saccade target selection deficits and compensation from pre-lesion to early and late post-lesion. (A) The proportion of correct saccades made to a single contralesional (yellow) or ipsilesional (light grey) target. (B) Point of equal selection on the free-choice paired stimulus trials. The point of equal selection is the stimulus onset asynchrony value at which the probability of choosing the contralesional (left) or ipsilesional (right) stimulus was equal. Positive y-axis values indicate that the point of equal selection was reached at a stimulus onset asynchrony in which the contralesional stimulus was presented before the ipsilesional stimulus, which would indicate a contralesional deficit. Negative y-axis values indicate a stimulus onset asynchrony in which the ipsilesional stimulus was presented first. Larger absolute values for the point of equal selection indicates more severe extinction-like deficits. (C) The proportion of saccades made to contralesional stimuli on trials with simultaneous presentation of both stimuli (stimulus onset asynchrony = 0 ms) on the free-choice trials. Statistical comparisons between pre-lesion and post-lesion time points were made using one-way ANOVAs with post-hoc Tukey’s tests (p < 0.05). Error bars represent standard error of the mean across sessions within each time point. Abbreviations: pre = pre-lesion; post1 = early post-lesion (week 1-2 post-lesion); post2 = late post-lesion (small lesion: week 8 post-lesion; large lesion: week 16 post-lesion); contra = contralesional; ipsi = ipsilesional.

### 2.4. Image acquisition at 7T

One hour prior to scanning, monkeys were sedated with intramuscular injections of 0.05-0.2 mg/kg acepromazine (Acevet 25 mg/ml) and 5.0-7.5 mg/kg ketamine (Vetalar 100 mg/ml), followed by 2.5 mg/kg propofol (10 mg/ml). Monkeys were then intubated with an endotracheal tube and anesthesia was maintained with 1.0-1.5% isoflurane mixed with 100% oxygen. Each monkey was positioned in a custom-built MRI primate bed with its head restrained to reduce motion and then inserted into the magnet bore for image acquisition, at which point the isoflurane level was maintained at 1.0% for the duration of the image acquisition. Body temperature, respiration, heart rate, and blood oxygen saturation levels were continuously monitored and were within a normal range throughout the scans. Body temperature was maintained using thermal insulation and a heating disk.

Imaging data were collected at pre-lesion (after behavioural training), week 1 post-lesion (early post-lesion), and at week 8 or 16 post-lesion when behaviour had compensated near pre-lesion baseline (late post-lesion). Although behaviour had compensated by week 8 for Monkey S, we were only able to obtain DWI data at week 16 post-lesion. Data were acquired on a 7T Siemens MAGNETOM Step 2.3 68-cm horizontal bore scanner (Erlangen, Germany) operating at a slew rate of 300 mT/m/s. We used an in-house designed and manufactured 8-channel transmit, 24-channel receive primate head radiofrequency coil for all image acquisitions (Gilbert et al., 2016). A high-resolution T2-weighted anatomical MR image was acquired using a turbo spin echo sequence with the following parameters: TR = 7500 ms, TE = 80 ms, slices = 42, matrix size = 320 × 320, field of view = 128 × 128 mm, acquisition voxel size = 0.4 mm x 0.4 mm x 1 mm. A T1-weighted MP2RAGE anatomical image was also acquired with these parameters: TR = 6500 ms, TE = 3.15 ms, TI1 = 800 ms, TI2 = 2700 ms, field of view = 128 × 128 mm, 0.5 mm isotropic resolution. Resting-state fMRI data were acquired and a detailed report of the fMRI acquisition has been previously published (Adam et al., 2020). In brief, we collected 4-6 10-minute runs of T2*-weighted continuous multi-band echo-planar imaging with 600 functional volumes per run using the following parameters: TR = 1000 ms, TE = 18 ms, slices = 42, slice thickness = 1 mm, and in-plane resolution = 1 mm x 1 mm.

DWI data were obtained using an interleaved echo planar imaging sequence with the following parameters: repetition time (TR) = 5100–7500 ms, echo time (TE) = 46.8–54.8 ms, number of averages = 1, number of slices = 46–54, slice thickness = 1 mm, in-plane resolution = 1 mm x 1 mm. We acquired 64 diffusion-encoding directions (b-value = 1000-1500 s/mm^2^) and one non-diffusion weighted volume (b-value = 0 s/mm^2^). Although there are slight within-subject variations in our TR (maximal difference of 1500 ms) and TE (maximal difference of 3 ms), previous work has shown no significant differences in the overall magnitude of diffusion between scans with larger differences in TR and TE (Celik, 2016). It has also been demonstrated that the mean FA in high angular resolution scans (e.g., 64 diffusion directions) was not significantly different between scans with a TR of 4000 ms or 13200 ms (Provenzale et al., 2018).

### 2.5. Image processing

Raw DWI data were converted from DICOM to NIFTI using MRIconvert (Lewis Center for Neuroimaging, University of Oregon) and reoriented to standard space using the FMRIB Software Library (FSL; http://www.fmrib.ox.ac.uk) tools ‘fslswapdim’ and ‘fslorient’. ASCII text files containing a list of gradient directions and b-values for each volume were also flipped and transposed to correspond with the reoriented DWI data. Data processing was then carried out using FMRIB’s Diffusion Toolbox (FDT) implemented with FSL. First, eddy current-induced distortions and subject motion were corrected using ‘EDDY’ (Andersson and Sotiropoulos, 2016). We then performed a diffusion tensor imaging (DTI) analysis to obtain four scalar maps representing FA and mean, axial, and radial diffusivity. DTI analysis involved fitting a tensor model at each voxel using ‘DTIFIT’ on the eddy corrected DWI data. The output DTI scalar maps are directly related to the three major eigenvalues (λ_1_, λ_2_, λ_3_) of the fitted tensor (i.e., the magnitude of diffusion for each eigenvector of the tensor). Axial diffusivity represents the magnitude of parallel diffusion and is defined as the first eigenvalue. Radial diffusivity represents the magnitude of perpendicular diffusion and is the average of the second and third eigenvalues. Mean diffusivity represents the total magnitude of diffusion and is the average of all three eigenvalues. Fractional anisotropy represents the degree of anisotropy and is calculated as the relative difference of the first eigenvalue compared to the other two eigenvalues (Alexander et al., 2007; Basser et al., 1994; Basser and Pierpaoli, 1996; Beaulieu, 2002).

Next, a multiple tensor model was fit at each voxel using ‘BEDPOSTX’ which estimates two fiber orientations per voxel to account for crossing fibers and more accurately generate probability distributions of local fiber orientations (Behrens et al., 2007, 2003b). This Bayesian estimation of multiple fiber directions vastly improves sensitivity when fiber tracking non-dominant pathways through regions of crossing fibers, such as the SLF (anterior-posterior) that has been traditionally difficult to track due to crossing dorsal-ventral projections in the more dominant corona radiata white matter bundle (Behrens et al., 2007). Transformation matrices were derived within subjects for each session from diffusion space to pre-lesion structural T2 space using a rigid-body transformation with 6 degrees of freedom using FSL’s ‘FLIRT’ (Jenkinson et al., 2002). The inverse transformation matrix from this registration was then used to register the seed masks from structural to diffusion space for the probabilistic tractography analysis. We have previously published preprocessing details for the resting-state fMRI data (Adam et al., 2020). Briefly, Resting-state fMRI data was processed using FSL and included brain extraction, MCFLIRT motion correction (6-parameter affine transformation), spatial smoothing (FWHM = 3 mm), high-pass temporal filtering, and registration to the standard macaque F99 template.

### 2.6. DWI tractography analysis

#### 2.6.1 Regions of interest for tractography

We reconstructed the contralesional and ipsilesional SLF and the transcallosal PFC and PPC tracts using bilateral seed regions (radius = 2 mm) created in pre-lesion structural T2 space for each subject. Seeds were placed in the frontal eye field (FEF) of the caudal PFC and in the lateral intraparietal area (LIP) of the PPC based on the (Saleem and Logothetis, 2006) rhesus monkey anatomical atlas. FEF seeds were placed in the anterior bank of the arcuate sulcus (Tehovnik et al., 2000) and LIP seeds were in the lateral bank of the intraparietal sulcus (Lewis and Van Essen, 2000). FEF and LIP constitute the main cortical nodes of the frontoparietal network (Wardak et al., 2011) and have been previously used to track these fibers in rhesus macaques (Hofer et al., 2008). Figure 4A shows representative seed mask locations in pre-lesion structural T2 space. The following seed region pairs were used in a probabilistic tractography analysis (described below) to reconstruct the four tracts of interest: bilateral FEF seeds were used to track the transcallosal PFC fiber tracts (Barbas and Pandya, 1984; Hofer et al., 2008); bilateral LIP seeds were used to track the transcallosal PPC tracts (Hofer et al., 2008); contralesional FEF and LIP seeds were used to track the contralesional SLF; and the ipsilesional FEF and LIP seeds were used to track the ipsilesional SLF (Petrides and Pandya, 1984; Schmahmann et al., 2007; Thiebaut de Schotten et al., 2012) (Fig. 4B). A midline sagittal exclusion mask was used to restrict tracking to the opposite hemisphere for the SLF association tracts and an axial exclusion mask at the anterior-posterior midpoint of the corpus callosum was used to restrict tracking to the anterior half of the brain for the transcallosal PFC tract or to the posterior half of the brain for the transcallosal PPC tract.

**Figure 4.**
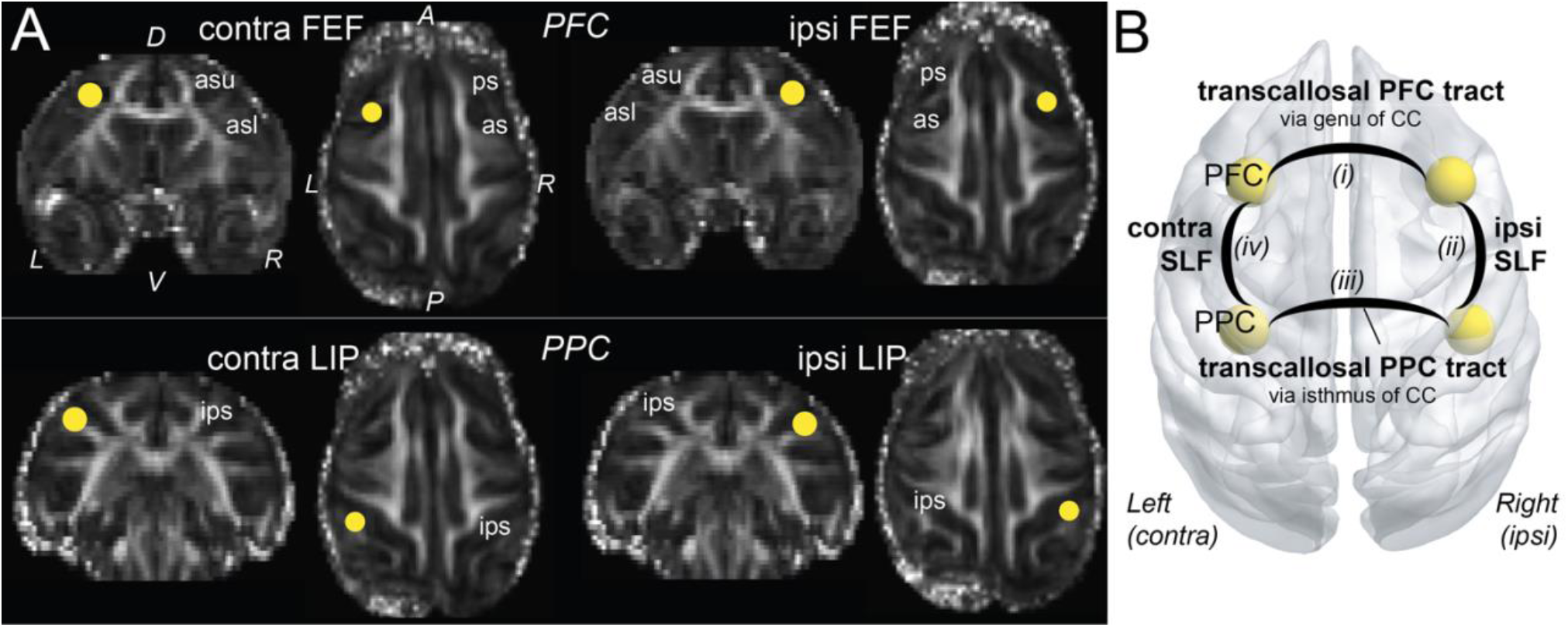
Seed masks to reconstruct fiber tracts of interest using probabilistic tractography. (A) Representative seed masks in bilateral FEF and LIP overlaid on FA maps in native T2 space shown at coronal and axial sections. Similar seeds were created for each monkey in native pre-lesion T2 space. (B) Schematic of the white matter tracts of interest. Probabilistic streamlines were generated for the (i) transcallosal PFC tract using bilateral FEF seeds, (ii) ipsilesional SLF association tract connecting the PFC and PPC using ipsilesional FEF and LIP seeds, (iii) transcallosal PPC tract using bilateral LIP seeds, and (iv) contralesional SLF association tract connecting the PFC and PPC using contralesional FEF and LIP seeds. Abbreviations: D = dorsal, V = ventral, L = left, R = right, contra = contralesional, ipsi = ipsilesional, A = anterior, P = posterior, as = arcuate sulcus, asu = upper limb of the arcuate sulcus, asl = lower limb of the arcuate sulcus, ips = intraparietal sulcus, ps = principal sulcus, FEF = frontal eye field, LIP = lateral intraparietal area, PFC = prefrontal cortex, PPC = posterior parietal cortex, SLF = superior longitudinal fasciculus, CC = corpus callosum.

#### 2.6.2 Probabilistic tractography

Probabilistic tractography was computed with FDT’s ‘ProbtrackX’ which uses the output from BEDPOSTX to estimate the number of streamlines that traveled between two seed regions for each voxel (Behrens et al., 2007, 2003a). We used the following ProbtrackX standard parameters: number of sample streamlines sent out per seed voxel = 5000, curvature threshold = 0.2, step length = 0.5, maximum number of steps = 2000, subsidiary fibre volume threshold = 0.01. Distance correction was additionally implemented to correct for the decrease in streamlines with distance from the seed region. For each of the 5000 streamlines per seed voxel sampled from the BEDPOSTX probability distribution, a ‘successful’ streamline was one that originated from one seed and reached the other. This algorithm outputs a streamline density map where individual voxel intensities represent the number of successful streamlines that passed through the voxel. The streamline density map was normalized by dividing it by the waytotal sum, which yielded voxel intensities that now represent the probability of that voxel being part of that tract. In contrast to methods that normalize streamline density maps using a constant proportion of the total number of streamlines sent out per voxel, proportional normalization based on the waytotal sum is the preferred approach when comparing reconstructed fiber tracts across sessions since it accounts for any differences in seeded voxels across sessions that may have affected trackability (Bennett et al., 2011). Streamline probability maps were then thresholded to maintain only voxels with intensities of at least 50% (i.e., a minimum of 50% probability that the voxel belongs to that streamline) and then visually inspected to confirm anatomical plausibility. Note that these suprathreshold white matter voxels are not necessarily exclusively part of the fiber tract of interest, but this probabilistic tractography approach gives a better approximation of the tract-related voxels compared to traditional approaches that use pre-defined region of interest FA mask without tractography. These normalization and thresholding procedures have been used for probabilistic tractography analysis (Cunningham et al., 2017; Galantucci et al., 2011; Gray et al., 2018; Latzman et al., 2015). Thresholded streamline probability maps representing the tracts of interest were generated for each subject and each session individually. These white matter fiber tracts have been identified in previous macaque studies using DWI tractography (Hofer et al., 2008; Hofer and Frahm, 2008; Schmahmann et al., 2007; Thiebaut de Schotten et al., 2012) and tracer methods (Barbas and Pandya, 1984; Petrides and Pandya, 1984). Figure 5 shows a representative sample of the reconstructed fiber tracts and the average streamline probability values for each tract are reported in Table 1.

**Table 1.** Average streamline probability of the suprathreshold voxels in the reconstructed white matter fiber tracts.

| Tract | Time since lesion | Monkey L |  |  | Monkey S |  |  | Monkey B |  |  | Monkey F |  |  |
| --- | --- | --- | --- | --- | --- | --- | --- | --- | --- | --- | --- | --- | --- |
|  |  | No. of voxels | Mean | SEM | No. of voxels | Mean | SEM | No. of voxels | Mean | SEM | No. of voxels | Mean | SEM |
| PFC–PFC | Pre | 888 | 87.3 | 0.56 | 641 | 88.6 | 0.66 | 1199 | 83.5 | 0.52 | 1211 | 84.6 | 0.50 |
|  | Post2 | 926 | 87.3 | 0.56 | 1126 | 87.1 | 0.51 | 1119 | 87.5 | 0.5 | 1151 | 82.4 | 0.56 |
| Ipsi SLF | Pre | 1949 | 81.4 | 0.41 | 801 | 82.6 | 0.66 | 1073 | 84.6 | 0.55 | 995 | 80.7 | 0.58 |
|  | Post2 | 930 | 82.2 | 0.61 | 1943 | 77.7 | 0.41 | 865 | 81.9 | 0.64 | 1498 | 75.1 | 0.48 |
| PPC–PPC | Pre | 1519 | 85.2 | 0.46 | 917 | 84.6 | 0.6 | 1653 | 85.9 | 0.43 | 1763 | 84.5 | 0.43 |
|  | Post2 | 1797 | 95.3 | 0.27 | 1141 | 85 | 0.55 | 1819 | 85.2 | 0.42 | 1604 | 84.3 | 0.46 |
| Contra SLF | Pre | 936 | 85.7 | 0.6 | 1072 | 85 | 0.55 | 1089 | 82.4 | 0.56 | 1421 | 80.4 | 0.49 |
|  | Post2 | 912 | 85.5 | 0.59 | 1047 | 85.3 | 0.55 | 1011 | 84 | 0.57 | 1421 | 78.8 | 0.49 |

All values were derived from thresholded probabilistic tracts with a minimum streamline probability of 50%. Abbreviations: No. = number; PFC = prefrontal cortex, PPC = posterior parietal cortex, SLF = superior longitudinal fasciculus, SEM = standard error of the mean, pre = pre-lesion, post2 = late post-lesion, ipsi = ipsilesional, contra = contralesional.

**Figure 5.**
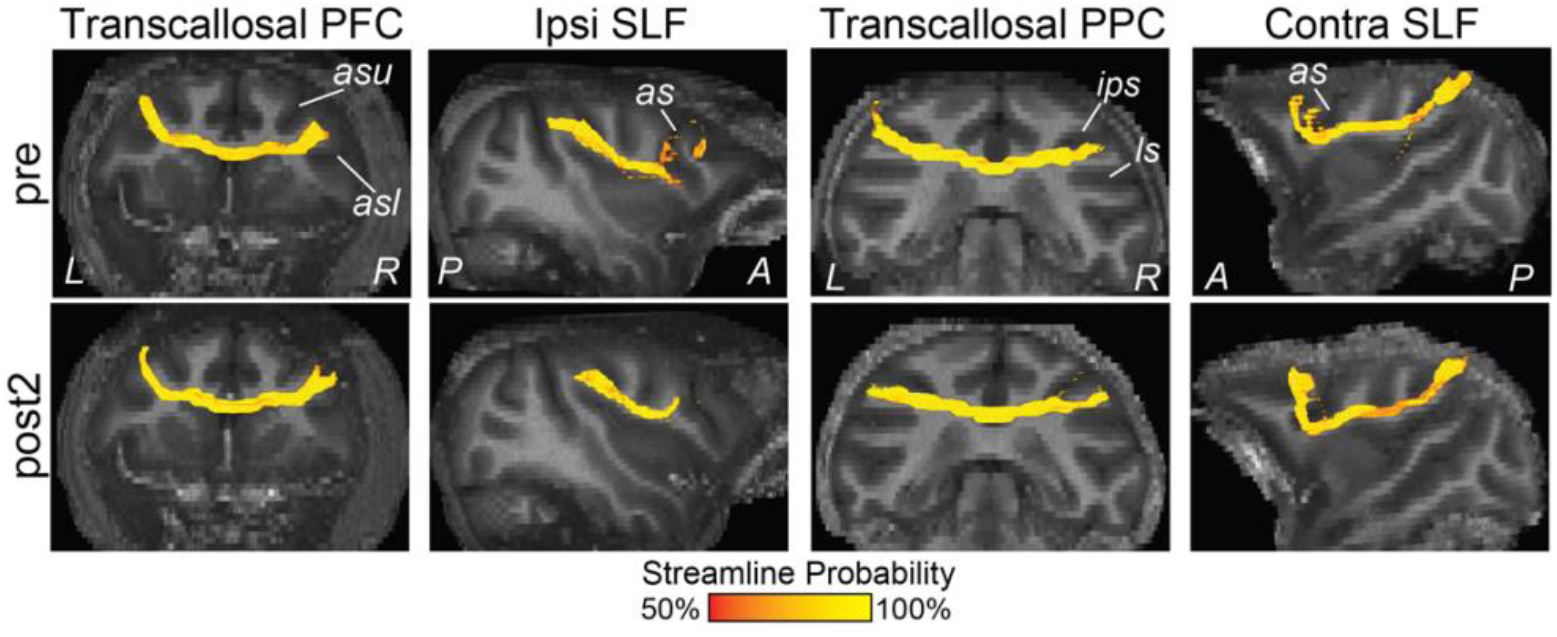
Representative white matter tracts reconstructed with probabilistic tractography. First column: bilateral FEF seeds revealed transcallosal streamlines between the bilateral PFC that traveled across hemispheres through the rostral portion, or genu, of the corpus callosum. Second column: ipsilesional FEF and LIP seeds revealed the ipsilesional SLF association fibers connecting frontal and parietal areas. Third column: bilateral LIP seeds revealed transcallosal streamlines between bilateral PPC that traveled across hemispheres through a posterior region (isthmus) of the corpus callosum. Fourth column: contralesional FEF and LIP seeds revealed the contralesional SLF association fibers connecting frontal and parietal areas. The colour bar represents streamline probabilities for each voxel in the thresholded tracts. Streamline probability maps are shown overlaid on a T2 coronal or parasagittal slice. Abbreviations: pre = pre-lesion, post2 = late post-lesion, A = anterior, P = posterior, L = left, R = right, PFC = prefrontal cortex, PPC = posterior parietal cortex, SLF = superior longitudinal fasciculus, contra = contralesional, ipsi = ipsilesional, as = arcuate sulcus, asu = upper limb of the arcuate sulcus, asl = lower limb of the arcuate sulcus, ips = intraparietal sulcus, ls = lateral sulcus.

### 2.7. Tract-specific DTI parameters

Here, we masked the four DTI scalar maps with the reconstructed tracts to obtain tract-specific measures of FA and mean, axial, and radial diffusivity at pre-lesion and late post-lesion. Previous studies have also obtained tract-specific measures of diffusivity and anisotropy since it takes fiber orientation into account, rather than only measuring diffusion parameters within pre-defined regions of interest without using masks generated from fiber tractography (Bennett et al., 2011; Galantucci et al., 2011; Ge et al., 2013; Gray et al., 2018; Lindenberg et al., 2012). First, the reconstructed thresholded tracts were binarized and only those voxels that overlapped in both the pre-lesion and late post-lesion binarized tracts were retained. This conservative approach accounts for any misalignment among individual tracts between time points. For the transcallosal tracts whose diffusion is largely oriented along the left-right x-direction (transcallosal PFC and PPC tracts), voxels within an x-coordinate range that were shared between pre-lesion and late pre-lesion tracts were retained. For the SLF association tracts whose diffusion is largely oriented along the anterior-posterior y-direction, voxels within a shared y-coordinate range in both pre-lesion and late post-lesion tracts were retained. Next, we masked DTI scalar maps with the binarized tracts to obtain tract-wise measures of FA and mean, axial, and radial diffusivity. We additionally calculated the average segment-wise FA values of three discrete segments along the length of each tract. Transcallosal PFC and PPC tracts were divided along the x-direction into contralesional/left, middle, and ipsilesional/right segments and SLF tracts were divided along the y-direction into anterior, middle, and posterior segments. This segment-wise spatial FA analysis may reveal important information about whether FA is uniform along the length of a tract and could identify whether local FA changes within one segment was driving changes in the average tract-wise FA (Colby et al., 2012; Davis et al., 2009). Tract-specific DTI metrics were statistically compared between pre-lesion and late post-lesion using two-sample t-tests (p < 0.05) with FDR correction for multiple comparisons within each subject.

### 2.8. Relationship between tract-wise FA and the rsFC of grey matter areas connected by that white matter fiber tract

We have previously reported the rsFC changes in these subjects along similar time points (Adam et al., 2020). Here, we compared changes in tract-wise FA with rsFC between cortical areas connected by that fiber tract. Bilateral seed regions of interest (radius = 2 mm) for the rsFC analysis were placed in areas of the resting-state frontoparietal network (Hutchison et al., 2012), including two major caudal PFC areas [FEF and 9/46D (DLPFC)] and nine PPC areas [PE, PEa, PEC, PF, PFG, POa (LIP), POaE, POal, Opt]. Caudal PFC areas are connected within hemisphere to the PPC areas via the SLF (Schmahmann et al., 2007). To compare tract-wise FA with rsFC, we extracted the average rsFC between groups of seed regions that corresponded to our tracts of interest: (1) contralesional and ipsilesional PFC seeds correspond with the transcallosal PFC tract, (2) ipsilesional PFC and PPC seeds correspond ipsilesional SLF, (3) contralesional and ipsilesional PPC seeds correspond with the transcallosal PPC tract, and (4) contralesional PFC and PPC seeds correspond with contralesional SLF. To obtain the rsFC between groups of seed regions, the average blood-oxygen level-dependent (BOLD) signal timecourse was first obtained for each seed and Pearson’s *r* correlation coefficients were computed between the BOLD signal timecourse of every pair of seeds, while regressing out the white matter and cerebrospinal fluid BOLD timecourse as noise. Fisher’s *r*-to-*z* transformation was applied to convert the correlation coefficients into z-scores, where z-scores denote the rsFC between seed regions. We averaged across the 4-6 z-score rsFC matrices for each session per subject, resulting in one rsFC matrix for pre-lesion and late post-lesion. For instance, to calculate the rsFC between contralesional PFC and PPC seed regions for comparison with the FA of the contralesional SLF, we took the |z-score| between the FEF and each of the nine PPC areas and the |z-score| between area 9/46D (DLPFC) and the nine PPC areas in the contralesional hemisphere, and used the average |z-score| as a rsFC index corresponding to the contralesional SLF. We statistically compared the |z-scores| (absolute rsFC) using two-sample t-tests with FDR correction for multiple comparisons (p < 0.05) from pre-lesion to late post-lesion.

## 3. Results

### 3.1. Longitudinal changes in the tract-specific DTI parameters

White matter tracts of interest were reconstructed using probabilistic tractography and used to extract tract-specific DTI parameters (FA and mean, axial, and radial diffusivity) from DTI scalar maps. Average values were calculated across the voxels of each tract within an overlapping x- or y-coordinate range between pre-lesion and late post-lesion. While we show results from all four tracts of interest, our main focus is on the remote fiber tracts that were not directly affected by the lesion, namely the contralesional SLF and transcallosal PPC tract.

In Monkey L, two-sample t-tests revealed that all four tracts showed significantly increased FA and decreased radial diffusivity from pre-lesion to late post-lesion, when behaviour had compensated (Fig. 6A,D). In addition, transcallosal PFC, contralesional SLF, and transcallosal PPC tracts had significantly decreased mean diffusivity, whereas the ipsilesional SLF had decreased axial diffusivity (Fig. 6B,C). For Monkey S, transcallosal PFC tract showed decreased FA whereas contralesional SLF and transcallosal PPC tracts showed increased FA (Fig. 6A). Both transcallosal PFC and ipsilesional SLF also showed increased mean, axial, and radial diffusivity, whereas transcallosal PPC and contralesional SLF showed decreased radial diffusivity (Fig. 6B–D). Lastly, contralesional SLF showed increased axial diffusivity (Fig. 6C). Findings shared by both small lesion monkeys were that the remote contralesional SLF and transcallosal PPC tracts showed increased FA and decreased radial diffusivity when behaviour had compensated.

**Figure 6.**
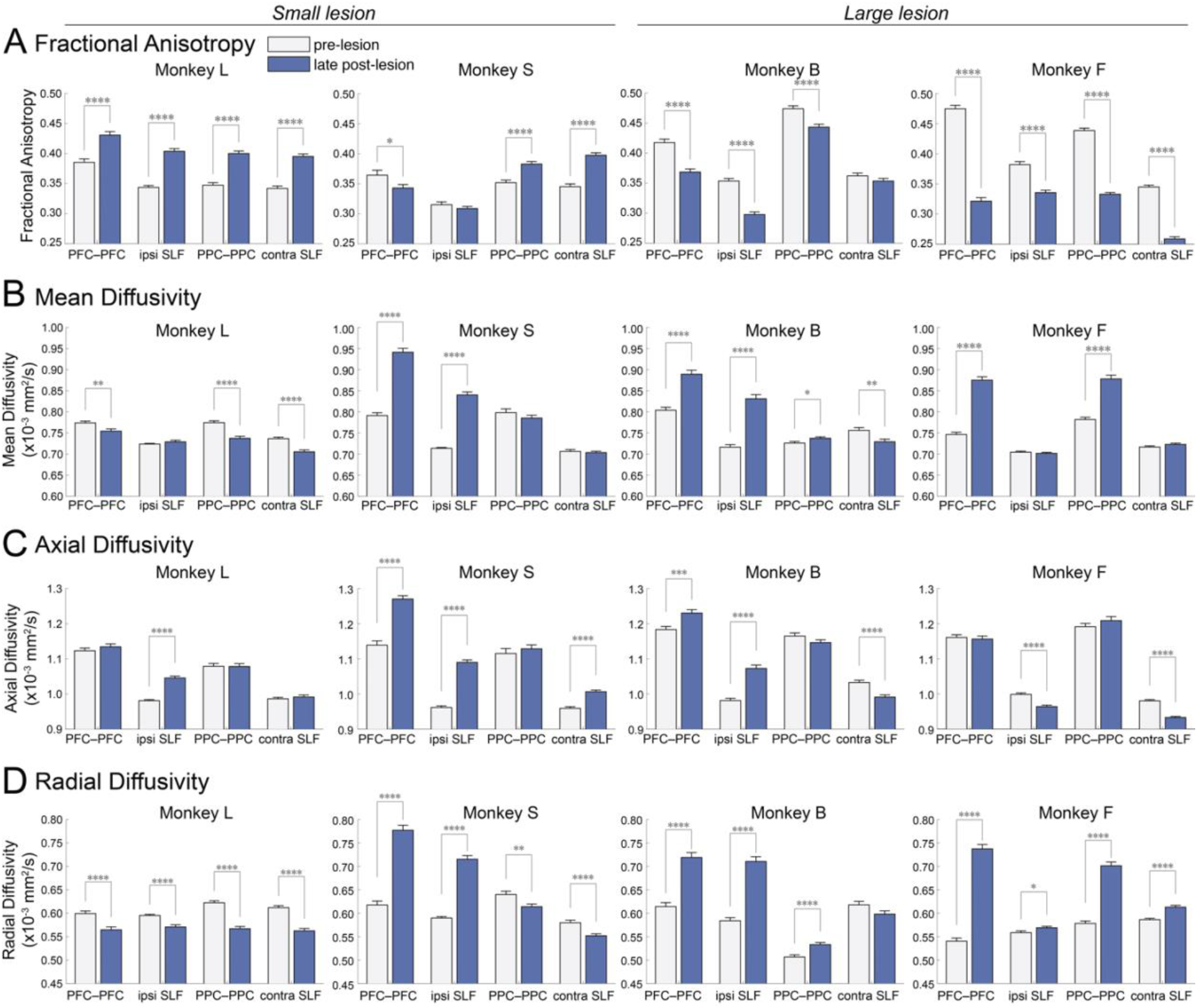
Changes in the average tract-wise DTI parameters over time. White matter tracts of interest were reconstructed with probabilistic tractography and then used to extract tract-specific measures of (A) fractional anisotropy, (B) mean diffusivity, (C) axial diffusivity, and (D) radial diffusivity. Statistical comparisons between pre-lesion and late post-lesion were made using two-sample t-tests with FDR correction for multiple comparisons (* = p<0.05, ** = p<0.01, *** = p<0.001, **** = p<0.0001). Error bars represent standard error of the mean across voxels. Abbreviations: pre = pre-lesion, post2 = late post-lesion (behavioural compensation time point), PFC–PFC = transcallosal PFC tract, PPC-PPC = transcallosal PPC tract, SLF = superior longitudinal fasciculus, ipsi = ipsilesional, contra = contralesional.

In Monkey B, transcallosal PFC and ipsilesional SLF showed significantly decreased FA and increased mean, axial, and radial diffusivity (Fig. 6A–D). Transcallosal PPC showed decreased FA and increased mean and radial diffusivity and lastly, contralesional SLF showed decreased mean and axial diffusivity. In Monkey F, decreased FA and increased radial diffusivity was found in all four tracts from pre-lesion to late post-lesion (Fig. 6A,D). In addition, transcallosal PFC and PPC tracts showed increased mean diffusivity (Fig. 6B), whereas contralesional and ipsilesional SLF showed decreased axial diffusivity (Fig. 6C). Findings shared by both large lesion monkeys were that (1) transcallosal PFC and transcallosal PPC showed decreased FA and increased mean and radial diffusivity, (2) ipsilesional SLF showed decreased FA and increased radial diffusivity, and (3) contralesional SLF showed decreased axial diffusivity.

### 3.2. Longitudinal changes of segment-wise FA in white matter fiber tracts

Here, we divided each tract into three segments to test whether FA changes were uniform along the length of the tract and to identify which segments likely drove the overall tract-wise FA. Transcallosal PFC and PPC tracts were divided along the x-direction into contralesional/left, middle, and ipsilesional/right segments and the SLF tracts were divided along the y-direction into anterior, middle, and posterior segments. Average FA was calculated for each segment and compared between pre-lesion and late post-lesion. In Monkey L, we found increased FA in the majority of segments across the four tracts (Fig. 7), except for the two segments closest to the lesion site (i.e., ipsilesional segment of the transcallosal PFC tract and anterior segment of the ipsilesional SLF) and the contralesional segment of the transcallosal PPC tract, which showed no change. In Monkey S, we found decreased FA in the middle segment of the transcallosal PFC tract and in the anterior segment of ipsilesional SLF (Fig. 7). In Monkey B, we found decreased segment-wise FA in the lesion-affected ipsilesional SLF and transcallosal PFC tracts and increased FA in the middle segment of the remote contralesional SLF and transcallosal PPC tracts (Fig. 7). In Monkey F, decreased FA was found in all tract segments, except for increased FA in the anterior segment of ipsilesional SLF (Fig. 7).

**Figure 7.**
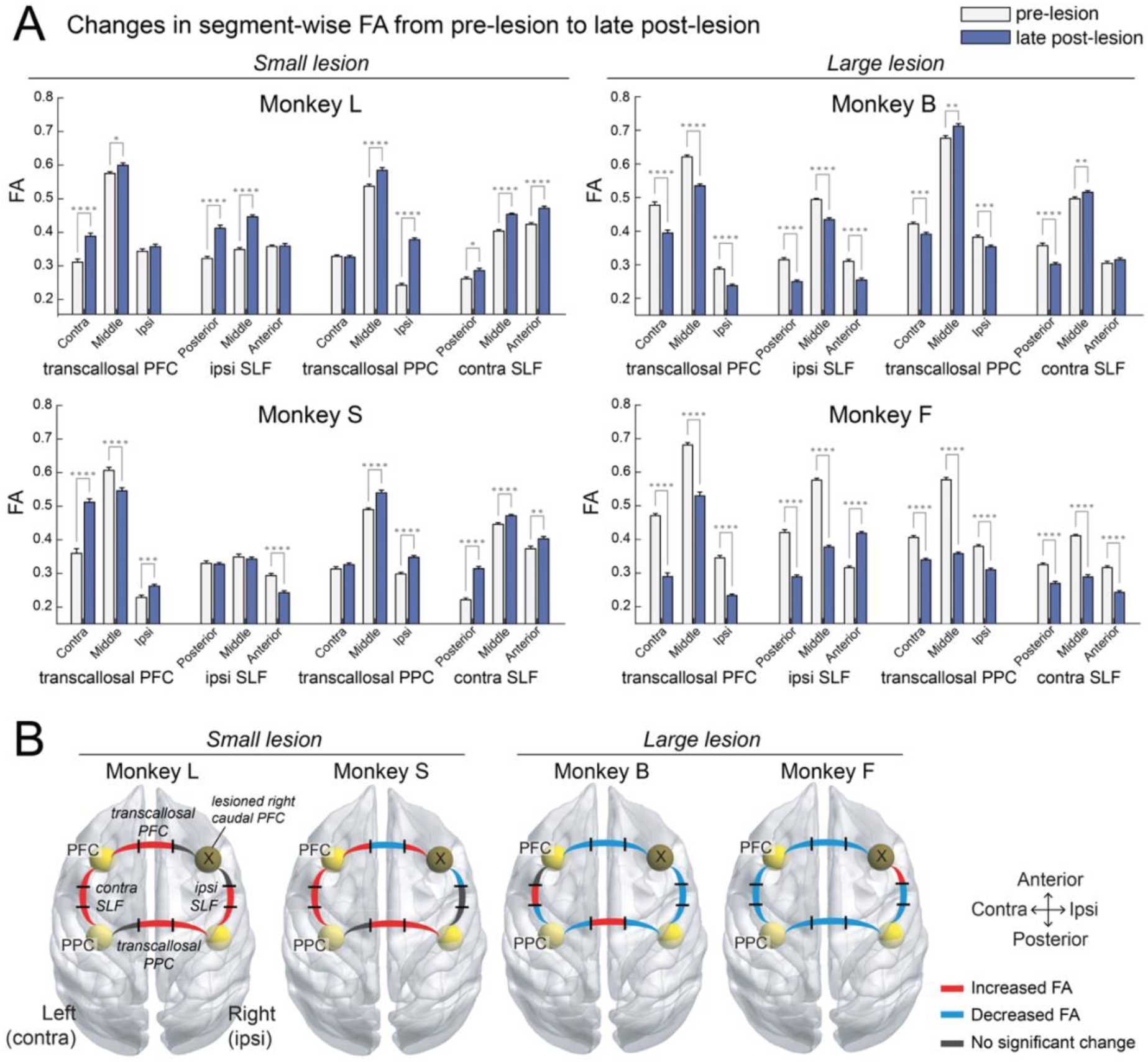
Changes in segment-wise FA of white matter tracts. (A) White matter tracts were divided along their length into three segments. Transcallosal PFC and PPC tracts were divided into contralesional/left, middle, and ipsilesional/right segments, and SLF tracts were divided into anterior, middle, and posterior segments. Average FA was extracted for each segment and statistically compared between pre-lesion and late post-lesion using two-sample t-tests with FDR correction for multiple comparisons (* = p<0.05, ** = p<0.01, *** = p<0.001, **** = p<0.0001). Error bars represent standard error of the mean across voxels. (B) Schematic summary of the segment-wise FA changes from pre-lesion to late post-lesion. Tract segments are illustrated with black lines dividing each segment. Red indicates significantly increased FA (p < 0.05), blue indicates significantly decreased FA (p < 0.05), and grey represents no significant change. Abbreviations: PFC = prefrontal cortex, SLF = superior longitudinal fasciculus, PPC = posterior parietal cortex, contra = contralesional side, ipsi = ipsilesional side.

### 3.3. Relationship between white matter tract microstructure and resting-state FC

We have previously published the changes in rsFC of the frontoparietal network in these subjects (Adam et al., 2020). Here, we examined the relationship between changes in the tract-wise FA with the corresponding change in rsFC (Table 2). We made the following rsFC–FA comparisons: (1) rsFC between contralesional PFC and PPC seeds with the FA of the contralesional SLF, (2) rsFC between ipsilesional PFC and PPC seeds with the FA of the ipsilesional SLF, (3) rsFC between bilateral PFC seeds with the FA of the transcallosal PFC tract, and (4) rsFC between bilateral PPC seeds with the FA of the transcallosal PPC tract.

**Table 2.** Changes in resting-state FC between frontoparietal areas of interest.

| Seed regions | Time | Monkey L |  |  | Monkey S |  |  | Monkey B |  |  | Monkey F |  |  |
| --- | --- | --- | --- | --- | --- | --- | --- | --- | --- | --- | --- | --- | --- |
|  |  | Mean | SEM | *p* | Mean | SEM | *p* | Mean | SEM | *p* | Mean | SEM | *p* |
| Bilateral PFC | Pre | 0.11 | 0.01 | 0.649 | 0.25 | 0.02 | 0.028* | 0.26 | 0.04 | 0.003* | 0.38 | 0.06 | 0.054 |
|  | Post2 | 0.10 | 0.01 |  | 0.18 | 0.01 |  | 0.43 | 0.02 |  | 0.26 | 0.02 |  |
| Ipsilesional PFC–PPC | Pre | 0.06 | 0.01 | 0.156 | 0.13 | 0.01 | 0.027* | 0.19 | 0.04 | 0.017* | 0.33 | 0.02 | 0.001* |
|  | Post2 | 0.08 | 0.01 |  | 0.08 | 0.01 |  | 0.30 | 0.02 |  | 0.21 | 0.01 |  |
| Bilateral PPC | Pre | 0.10 | 0.01 | 0.997 | 0.19 | 0.03 | 0.492 | 0.27 | 0.04 | 0.019* | 0.51 | 0.03 | 0.873 |
|  | Post2 | 0.10 | 0.01 |  | 0.17 | 0.00 |  | 0.38 | 0.02 |  | 0.50 | 0.02 |  |
| Contralesional PFC–PPC | Pre | 0.06 | 0.01 | 0.001* | 0.13 | 0.02 | 0.048* | 0.20 | 0.04 | 0.001* | 0.40 | 0.05 | 0.644 |
|  | Post2 | 0.12 | 0.01 |  | 0.19 | 0.01 |  | 0.47 | 0.03 |  | 0.42 | 0.02 |  |

Abbreviations: pre = pre-lesion, post2 = late post-lesion (behavioural recovery time point),
contra = contralesional, ipsi = ipsilesional, SEM = standard error of the mean, *p* = significance
value between pre- and post-lesion, \* = *p* < 0.05.

The rsFC between contralesional PFC–PPC (corresponding to contralesional SLF) was increased from pre-to late post-lesion across the four monkeys, except this effect was not significant for Monkey F (Table 2). In the two small lesion monkeys, Monkey L and Monkey S, this increased rsFC corresponded with the increased tract-wise FA in the contralesional SLF (Fig. 8). However, the two large lesion monkeys (Monkeys B and F) did not show a corresponding FA increase in contralesional SLF; instead FA had decreased in both monkeys, but this effect was not significant for Monkey B (see Fig. 6). Monkey S additionally showed decreased rsFC between bilateral PFC, which corresponded with decreased FA in the transcallosal PFC tract (Fig. 8), and decreased rsFC in ipsilesional SLF (Table 2). In Monkey B, increased rsFC was found in all comparisons which were inconsistent with the decreased FA found in those tracts (Fig. 8). Monkey F showed decreased rsFC between bilateral PFC which corresponded with decreased FA in the transcallosal PFC tract. Overall, the direction of rsFC changes largely matched the changes in tract-wise FA in the small lesion monkeys, but not in the large lesion monkeys.

**Figure 8.**
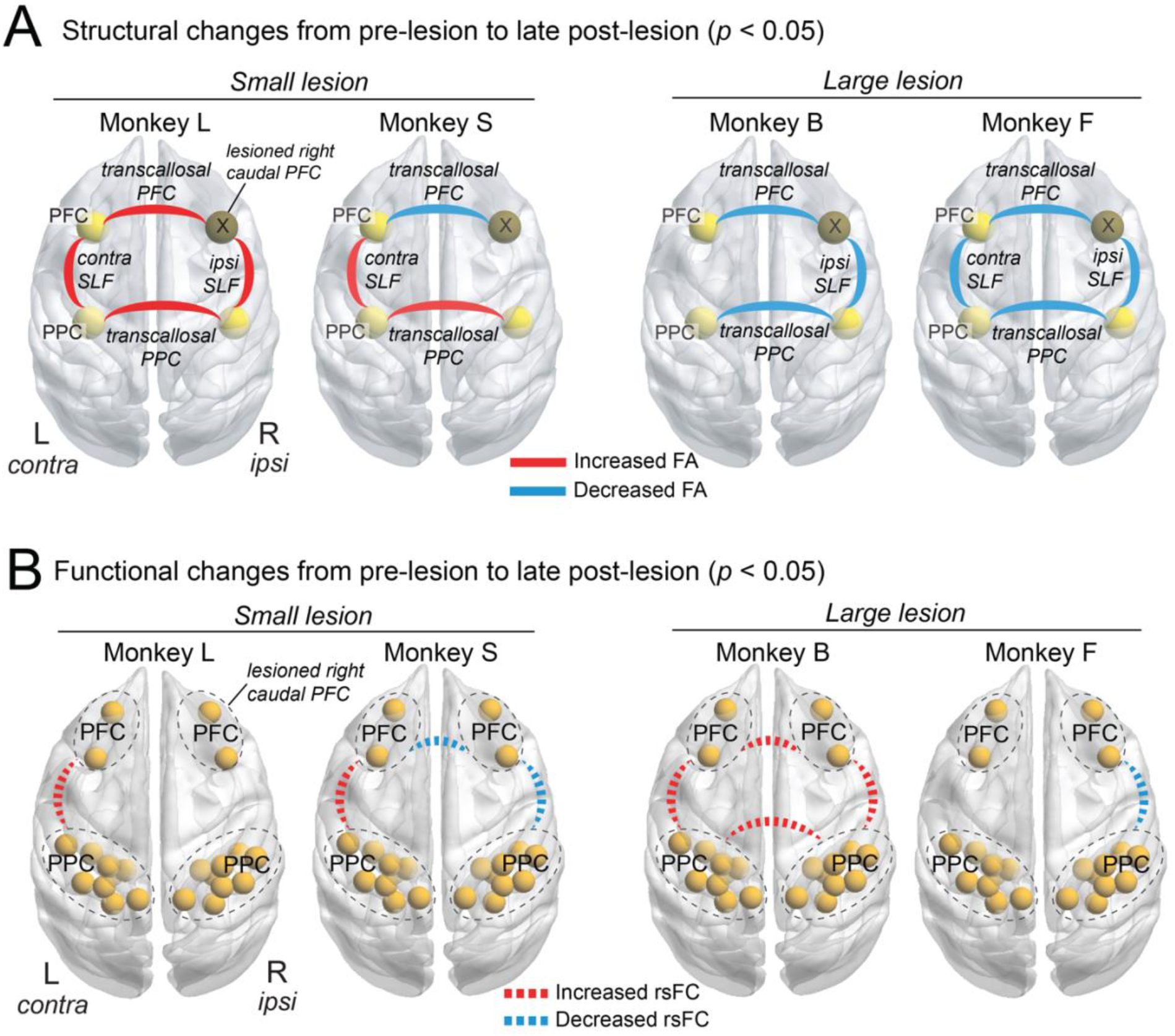
Schematic summary of the changes in tract-specific FA and rsFC. (A) Changes in the tract-wise mean FA for each white matter tract of interest from pre-lesion to late post-lesion, when behavioural performance had compensated. Solid red lines indicate significantly increased FA and solid blue lines indicate significantly decreased FA. (B) Resting-state FC changes that correspond to the white matter tracts of interest. Resting-state FC was calculated as the average absolute z-score between all pairwise seed regions and compared with the FA of the corresponding white matter tract. Dotted red lines indicate significantly increased FC and dotted blue lines indicate significantly decreased FC. Statistical comparisons between pre-lesion and late post-lesion were made using two-sample t-tests with FDR correction for multiple comparisons (p < 0.05). Abbreviations: PFC = prefrontal cortex, SLF = superior longitudinal fasciculus, PPC = posterior parietal cortex, contra = contralesional, ipsi = ipsilesional.

## 4. Discussion

In this longitudinal DWI study, we used probabilistic tractography to investigate microstructural changes in frontoparietal white matter tracts after right caudal PFC lesions in macaque monkeys. DTI metrics were obtained within each tract and compared from pre-lesion to late post-lesion, when behavioural performance on a saccade choice task had largely recovered. We have previously published detailed reports of the behaviour and resting-state fMRI data in these subjects (Adam et al., 2020, 2019). Here, we found that tract-wise FA in remote contralesional SLF and transcallosal PPC tract was differentially altered based on lesion size, with increased FA after small PFC lesions (Monkeys L and S) and decreased FA after larger lesion (Monkeys B and F). This study also highlights the importance of evaluating segment-wise FA since the changes in FA were not always uniform along the length of a fiber tract. The lack of consistent or compensatory changes in network-wide rsFC and FA after larger lesions may suggest the recruitment of alternate pathways beyond the cortical frontoparietal network to support the behavioural recovery.

### 4.1. White matter degeneration in the lesion-affected white matter tracts

White matter alterations after a focal lesion initially occur locally in perilesional tissue and along fiber tracts directly connected to the site of the lesion by anterograde (i.e., Wallerian) and retrograde axonal degeneration (Beaulieu, 2002; Pierpaoli et al., 2001; Thomalla et al., 2004; Werring et al., 2000). These changes in perilesional tissue microstructure have been studied using measures of FA from DWI studies in stroke patients (Pierpaoli et al., 2001; Thomalla et al., 2004; Umarova et al., 2017; Werring et al., 2000). In this section, we discuss the microstructural changes in the lesion-affected ipsilesional SLF and transcallosal PFC tracts (i.e. tracts that directly innervate the lesioned right caudal PFC).

Ipsilesional SLF and transcallosal PFC tracts in Monkey S, Monkey B, and Monkey F show decreased FA and increased radial diffusivity. DTI studies in stroke patients have also reported decreased FA and increased radial diffusivity in lesion-affected white matter tracts (Dacosta-Aguayo et al., 2014; Schaechter et al., 2009; Umarova et al., 2017). Previous reports of white matter degeneration on DWI suggest that this pattern of microstructural changes reflects myelin breakdown in axons directly connected to the lesion site (Beaulieu, 2002; Pierpaoli et al., 2001; Werring et al., 2000). In Monkey S, segment-wise FA analysis of the transcallosal PFC tract revealed that only the middle segment showed a decreased FA, which likely drove the decreased tract-wise FA for that tract. This may be due to FA changes within other prefrontal fibers traversing the genu of the corpus callosum that were not picked up by our tractography analysis and may have been more impacted by the lesion.

In contrast, Monkey L showed increased FA in ipsilesional SLF and transcallosal PFC tracts at late post-lesion. Since Monkey L sustained the smallest and most focal lesion, it is possible that there was a relatively greater number of preserved/undamaged axonal fibers from the lesioned caudal PFC traveling within hemisphere via ipsilesional SLF or between hemispheres via transcallosal PFC fibers. Spared fibers may have allowed for optimal neural compensation strategies to take place by way of local plasticity in the perilesional tracts (Murphy and Corbett, 2009). While increased FA in ipsilesional SLF and transcallosal PFC tracts in Monkey L may be viewed as the outcome of adaptive plasticity for behavioural recovery, these findings should be interpreted with caution since DWI changes in lesion-affected tracts may be confounded by direct lesion pathology (Pierpaoli et al., 2001).

### 4.2. Transneuronal degeneration or compensation in the remote white matter tracts

Over several days to weeks following the initial degeneration of fibers directly connected to the lesion, white matter atrophy may take place in remote areas indirectly connected to the lesion. Fibers connected to the lesion across multiple synapses may undergo anterograde and retrograde transneuronal degeneration (Baron et al., 2014; Fornito et al., 2015; Grayson et al., 2017; Zhang et al., 2012), the extent of which likely depends on the lesion size and location (Thiel et al., 2010; Wasserman and Schlichter, 2008). Here, we focus on the remote fiber tracts that are indirectly connected to the lesioned right caudal PFC, namely the contralesional SLF and transcallosal PPC tracts. Transneuronal degeneration may appear on DWI as decreased FA, decreased axial diffusivity, and/or increased radial diffusivity in white matter tracts remote from the lesion (Beaulieu, 2002). These diffusion changes may reflect a combination of degenerative changes, including decreased fiber density, demyelination, and axonal loss (Sotak, 2002).

We found evidence of transneuronal degeneration in the contralesional SLF and transcallosal PPC tracts in the two large lesion monkeys, but not in the small lesion monkeys. Specifically, Monkey B showed decreased FA and increased radial diffusivity in transcallosal PPC and decreased axial diffusivity in contralesional SLF. Monkey F showed decreased FA and increased radial diffusivity in both tracts and additionally decreased axial diffusivity in contralesional SLF. Several lines of evidence support our finding of decreased FA in remote fiber tracts only after larger lesions and more severe/lasting deficits. DWI studies have reported that, compared to patients with mild or recovered neglect, patients with severe or persistent deficits showed decreased FA in the posterior corpus callosum, which provides the communication link between parietal areas in the damaged and intact hemispheres (Bozzali et al., 2012; Lunven et al., 2015), or decreased FA between contralesional frontoparietal areas (Umarova et al., 2014). Similarly, in a longitudinal study of neglect, Umarova et al. (2017) reported that the degree of unrecovered neglect correlated strongly with white matter degeneration in the intact hemisphere between contralesional frontoparietal pathways (Umarova et al., 2017). This finding has also been demonstrated in one other study in chronic stroke patients recovering from motor-related deficits, such that poorly recovered patients had reduced FA in both ipsilesional and contralesional corticospinal tracts, whereas well-recovered patients showed increased FA in those tracts compared to healthy controls (Schaechter et al., 2009). Alternatively, decreased FA in remote tracts after larger lesions may not necessarily underlie the severity or persistence of deficits, but may instead reflect an epiphenomenon of larger lesions (Umarova et al., 2017). It is possible that the lesions in Monkeys L and S were not large enough to induce transneuronal degeneration. In addition, the segment-wise FA analysis in Monkey B showed increased FA in the middle segments of contralesional SLF and transcallosal PPC tracts that were averaged out in the tract-wise mean FA. This increased FA may represent an adaptive change that supports behavioural recovery, since Monkey B showed a greater degree of recovery than Monkey F. However, It would be valuable for future studies to test whether these spatial differences in FA along fiber tracts are due to true microstructural alterations or reflect artifacts from crossing fibers (Jones et al., 2013).

In contrast, the two small lesion monkeys (Monkeys L and S) showed increased FA and decreased radial diffusivity in the remote contralesional SLF and transcallosal PPC tracts. Increased FA combined with decreased radial diffusivity likely reflects increased myelination of these remote fiber tracts (Beaulieu, 2002). It is also possible that this increased FA in the remote fiber tracts after smaller lesions reflects an adaptive or compensatory change in white matter microstructure that may be related to the faster time to behavioural recovery in these animals (8 weeks) compared to those with larger lesions (16 weeks). Previous studies have highlighted an adaptive role for post-lesion changes in distant white matter tissue in both stroke patients (Bütefisch et al., 2003; Crofts et al., 2011; Lin et al., 2015; Liu et al., 2015; Schaechter et al., 2009) and animal models (Carmichael and Chesselet, 2002; Liu et al., 2008; Napieralski et al., 1996; Stroemer et al., 1995). Specifically, improved motor function in stroke patients correlated with increased FA in contralesional white matter (Lin et al., 2015; Liu et al., 2015; Schaechter et al., 2009).

One possibility is that the increased FA after small lesions and decreased FA after larger lesions in remote fiber tracts may reflect differences in the extent of disinhibition and potential downstream excitotoxicity. Focal lesions can lead to large-scale depolarization of connected areas resulting in disinhibition and hyperexcitability of widespread, functionally related networks (Buchkremer-Ratzmann and Witte, 1997; Fornito et al., 2015; Liepert et al., 2000). Adaptive structural plasticity after a focal lesion may be induced by this hyperexcitability, which has been associated with axonal and dendritic growth of undamaged fibers, myelin remodeling, synaptogenesis (Carmichael and Chesselet, 2002; Fornito et al., 2015; Gonzalez and Kolb, 2003; Jones and Schallert, 1992; Lin et al., 2015) and improved motor function (Reitmeir et al., 2011). However, larger lesions induce more widespread disinhibition and may lead to remote white matter degeneration across connected areas due to excitotoxicity and excessive metabolic stress from persistent hyperactivation (Buchkremer-Ratzmann and Witte, 1997; W. de Haan et al., 2012; Fornito et al., 2015; Ross and Ebner, 1990; Saxena and Caroni, 2011). Smaller lesions in Monkeys L and S may not have been sufficient enough to induce maladaptive hyperactivation in remote areas; here, the degree of disinhibition/hyperexcitability may have allowed for adaptive plasticity and contributed to compensatory changes across the functionally related network. On the other hand, larger lesions in Monkeys B and F likely induced more substantial disinhibition across the bilateral network and subsequently led to excessive hyperactivation, excitotoxicity, and metabolic stress, ultimately resulting in transneuronal degeneration of remote fiber tracts.

### 4.3. Relationship between white matter microstructure and resting-state FC

Although we found inconsistencies between DTI-derived metrics and resting-state FC, the magnitude of significant change reported for FA and rsFC are very robust. Thus, these discrepancies are unlikely to result from methodology, but instead may reflect variability in the compensatory response to lesions of different size or affecting different areas. Increased FA in the remote contralesional SLF and increased rsFC between corresponding grey matter areas (contralesional PFC–PPC) in both small lesion monkeys was the only consistent finding between FA and rsFC. We interpret this paired increase in contralesional FA and rsFC as support for a compensatory role of the contralesional hemisphere in the recovery of function after small PFC lesions. In our previous resting-state fMRI study, we reported that rsFC between areas in the contralesional PFC and ipsilesional PPC correlated with improving behavioural performance over time in all four monkeys (Adam et al., 2020). Since the contralesional SLF is one of the pathways that contributes to the indirect link between contralesional PFC and ipsilesional PPC, it is possible that the increased FA in contralesional SLF mediated the increased rsFC between contralesional PFC and ipsilesional PPC areas. This interpretation is supported by previous studies that showed positive correlations between rsFC and structural connectivity/FA in the white matter tracts that contribute to the indirect/polysynaptic pathway linking the functionally connected areas (Adachi et al., 2012; Greicius et al., 2009; Honey et al., 2009; Hori et al., 2020; Messé et al., 2014).

However, this compensatory response was not observed in the two large lesion monkeys. In Monkey B, rsFC increased between all areas from pre-lesion to late post-lesion, yet this was in contrast to the significantly decreased FA in ipsilesional SLF, transcallosal PFC, and transcallosal PPC tracts. Notably, there was no significant change in tract-wise FA for contralesional SLF even though the corresponding rsFC was significantly increased. One interpretation is that although functional and structural connectivity are often correlated, increased functional connectivity between two regions with a ‘weakened’ structural connection (e.g., decreased FA) may be mediated by strengthening of structural connections with a related third region (Honey et al., 2009; Koch et al., 2002). Another possible explanation for the opposite change in FA and rsFC in Monkey B comes from accumulating evidence demonstrating that network reorganization can maintain functional connectivity after loss of major structural pathways (O’Reilly et al., 2013; Tyszka et al., 2011; Uddin, 2013; Uddin et al., 2008). After major disconnections of the corpus callosum, sparing of even a few commissural fibers was sufficient to maintain normal levels of functional connectivity between hemispheres months later (O’Reilly et al., 2013; Tyszka et al., 2011; Uddin, 2013; Uddin et al., 2008). On the other hand, Monkey F did not show similar significant widespread increases in rsFC as in Monkey B, but instead only had significantly decreased rsFC in ipsilesional SLF. The lack of any significantly increased rsFC along with an overall decreased FA in all fiber tracts in Monkey F at the time of behavioural recovery suggests that neural compensation may have involved other brain areas or networks. Altogether, decreased FA in frontoparietal white matter tracts in both large lesion monkeys hint that behavioural recovery after larger PFC lesions may not be solely mediated by cortical connections in the frontoparietal network. Instead, there may be a functionally relevant third region/network that supports behavioural compensation and possibly works to maintain cortical rsFC (Damoiseaux and Greicius, 2009).

### 4.5. Conclusions

After the recovery of contralesional saccade choice deficits, FA in remote contralesional SLF and transcallosal PPC tracts was increased in monkeys with small PFC lesions, and decreased in monkeys with larger lesions compared to pre-lesion. This suggests that the white matter tracts connecting remote areas of the frontoparietal network (i.e., distant to the lesion) may contribute an important compensatory response to support recovery of function after small PFC lesions. However, larger lesions may have induced more widespread damage to the structural network over time such that these remote fiber tracts are no longer sufficiently able to compensate for lost function. Future research is needed to clarify the behavioural relevance of the remote fiber tracts and to investigate an alternate source of neural compensation after greater frontoparietal network damage.

## Acknowledgements

We thank Joseph Gati and Trevor Szekeres for technical support and assistance with MRI acquisition; Nicole Hague and Ashley Kirley for animal care and preparation; and Kyle Gilbert for designing the macaque RF coil.

